# Environmental transcriptomics under heat stress: Can environmental RNA reveal changes in gene expression of aquatic organisms?

**DOI:** 10.1101/2022.10.06.510878

**Authors:** Robert M. Hechler, Matthew C. Yates, Frédéric J. J. Chain, Melania E. Cristescu

## Abstract

To safeguard biodiversity in a changing climate, we require taxonomic information about species turnover and insights into the health of organisms. Environmental DNA approaches are increasingly used for species identification, but cannot provide functional insights. Transcriptomic methods reveal the physiological states of macroorganisms, but are currently species specific and require tissue sampling or animal sacrifice, making community-wide assessments challenging. Here, we test if broad functional information (expression level of the transcribed genes) can be harnessed from environmental RNA (eRNA), which includes extra-organismal RNA from macroorganisms along with whole microorganisms. We exposed *Daphnia pulex* as well as phytoplankton prey and microorganism colonizers to control (20 °C) and heat stress (28 °C) conditions for seven days. We sequenced eRNA from tank water (after complete removal of *Daphnia*) as well as RNA from *Daphnia* tissue, enabling comparisons of extra-organismal and organismal RNA based gene expression profiles. Both RNA types detected similar heat stress responses of *Daphnia*. Using eRNA, we identified 32 *Daphnia* genes to be differentially expressed following heat stress. Of these, 17 were also differentially expressed and exhibited similar levels of relative expression in organismal RNA. In addition to the extra-organismal *Daphnia* response, eRNA detected community-wide heat stress responses consisting of distinct functional profiles and 121 differentially expressed genes across 8 taxa. Our study demonstrates that environmental transcriptomics based on eRNA can non-invasively reveal gene expression responses of macroorganisms following environmental changes, with broad potential implications for the biomonitoring of ecological health across the trophic chain.

## Introduction

In an era of ecological crises, monitoring the presence and health of organisms within an ecosystem is essential for the preservation of biodiversity. Environmental DNA is routinely used for biomonitoring, but such applications are limited to species detection (Cristescu & Hebert, 2018; Deiner et al., 2017). The RNA molecule provides an additional layer of functional information, as transcribed genes reflect an organism’s physiological status (Huang et al., 2002). Thus, transcriptomics is used to infer the ecological health of organisms, but such surveys are species-specific and dependent on sampling organismal RNA (oRNA) directly, which is labor-intensive and requires tissue collection or animal sacrifice (Alvarez et al., 2015; Baillon et al., 2015; Gleason & Burton, 2015; Houde et al., 2019; Jeffries et al., 2021; Miller et al., 2017). Non-invasive environmental RNA (eRNA) approaches could overcome these limitations by providing insights into the health of populations and communities (Amarasiri et al., 2021; Cristescu, 2019; Veilleux et al., 2021; Yates et al., 2021). However, using eRNA to inform on ecological health has only been theorized (see Yates et al., 2021) and remains yet to be empirically tested.

We broadly define eRNA as RNA extracted from the environment, including extra-organismal RNA released by macroorganisms as well as whole microorganisms. Microbiologists have long extracted RNA from bulk samples to profile gene expression in microbial communities, referred to as metatranscriptomics (Frias-Lopez et al., 2008; Gilbert et al., 2008; Poretsky et al., 2005). Despite more than two decades of research, metatranscriptomics remains limited to studying the gene expression of microorganisms captured in bulk samples rich in oRNA. The general assumption has been that the eRNA released by macroorganisms into the environment is labile and degrades too rapidly to be reliably detected or quantified (but see Cristescu, 2019). However, the RNA shed by macroorganisms into their surrounding environment can be successfully extracted and quantified (T. Jo et al., 2022; Kagzi et al., 2022; Littlefair et al., 2022; Marshall et al., 2021; Miyata et al., 2021; Tsuri et al., 2021; von Ammon et al., 2019; Wood et al., 2020). Despite this evidence supporting the extractability of eRNA, environmental transcriptomics, defined here as eRNA based gene expression profiling, remained untested.

In aquatic systems, environmental conditions modulate biodiversity and are reflected in changes of metatranscriptomes (Frias-Lopez et al., 2008; Salazar et al., 2019; Sunagawa et al., 2015; Sunday et al., 2011). Temperature is a key abiotic factor that influences the physiology and fitness of ectotherms due to the relationship between body and environmental temperatures (Huey & Berrigan, 2001; Schulte, 2015; Rodgers, 2021). Rapid increases in temperature, such as heatwaves, can exceed the thermal limits of aquatic ectotherms and exponentially increase standard metabolic rates, resulting in decreases in fitness and survival (Harley et al., 2006; Reid et al., 2019; Rodgers, 2021; Schulte, 2015). Transcriptomics is widely used to study the heat stress response of many taxa, including fish (Akbarzadeh et al., 2018; Houde et al., 2019; Narum & Campbell, 2015; Veilleux et al., 2015, 2018), coral holobionts (Savary et al., 2021; Voolstra et al., 2021), molluscs (Chen et al., 2019; Gleason & Burton, 2015), and copepods (Kelly et al., 2017; Semmouri et al., 2019). *Daphnia pulex*, a key bioindicator species, exhibits widespread downregulation of genes involved in metabolic processes under heat stress (Becker et al., 2018; Yampolsky et al., 2014). Clearly, gene expression surveys can provide species-specific functional insights important for conserving aquatic biodiversity (Bozinovic & Pörtner, 2015; Evans & Hofmann, 2012; Wikelski & Cooke, 2006). However, a species by species analysis based on oRNA cannot capture community-wide changes without extensive efforts. Mapping eRNA reads to the reference genomes of multiple target species and using existing metatranscriptomic pipelines, could potentially enable environmental transcriptomics to reveal gene expression responses across the trophic chain.

We constructed simple mock freshwater communities containing *Daphnia pulex*, as well as three phytoplankton species and opportunistic microorganisms that colonized the artificial lake water. The communities were exposed to control (20 °C) and near lethal *Daphnia* heat stress (28 °C) conditions for seven days. To enable comparisons between eRNA and oRNA based gene expression profiles, we sequenced eRNA from tank water (after complete *Daphnia* removal) and oRNA from *Daphnia* tissue. We conducted differential gene expression analyses between the control and heat stress conditions and compared the *Daphnia* functional stress response identified in eRNA and oRNA. In addition to the *Daphnia* analysis, we used eRNA to investigate the community-wide heat stress response. We hypothesized that 1) eRNA captures transcriptional responses to heat stress and 2) eRNA contains a subset of the differentially expressed *Daphnia* genes observed in oRNA after exposure to heat stress. Based on our findings, we discuss the potential of eRNA based transcriptomics for the biological monitoring of ecological communities and also provide recommendations to overcome potential limitations.

## Materials and Methods

### Experimental design

We constructed simple mock freshwater communities composed of *Daphnia pulex*, three algae species (*Ankistrodesmus falcatus, Scenedesmus quadricauda*, and *Pseudokirchneriella subcapitata*) and microorganisms that colonized the artificial lake water. The communities were exposed to 20 °C (control) and 28 °C (neat-lethal stress for Daphnia) temperatures for seven days to mimic natural heatwave conditions and allow for eRNA accumulation in tank water. Communities were reared in 8 L tanks containing artificial lake water (Celis-Salgado et al., 2008) and were seeded with a starting population of 50 parthenogenetic females of *Daphnia pulex* (clone from Illinois, USA; 40.24, −87.78). *Daphnia* populations were kept in controlled chambers for 45 days at control growing conditions: 20 °C, relative humidity of 50% and a photoperiod of 16:8 hour light:dark. *Daphnia* were fed 1.5 mL algae (1:1:1 ratio of *A. falcatus, S. quadricauda*, and *P. subcapitata*) twice per week. After 45 days, the *Daphnia* populations in all tanks reached a minimum of 500 individuals and we began the experiment by transferring tanks to controlled experimental chambers pre-set to the respective temperatures (20 °C and 28 °C).

### Sample collection

Each temperature treatment had four tanks (biological replicates), and we collected four eRNA and oRNA technical replicates per tank. We stirred the tanks for 60 seconds and filtered 500 mL of water through a 60 μm mesh to remove all *Daphnia* and ephippia. To capture eRNA, the *Daphnia* free water was then filtered through a 0.7 μm glass microfiber filter. Each filter was cut in half and placed in individual 1.5 mL microcentrifuge tubes containing 370 μL RLT buffer (Qiagen) and 3.7 μL β-mercaptoethanol. Tubes were immediately stored at −80 °C until eRNA extraction. Negative eRNA filtration control samples (500 mL of distilled water) were collected at the beginning, middle and end of the sampling day, using the same methods. For each oRNA sample, six live organisms (3 adults and 3 juveniles) per tank were haphazardly collected using a sterile pipette and transferred to a 1.5 mL microcentrifuge tube containing 370 μL RLT buffer (Qiagen) and 3.7 μL β-mercaptoethanol. To maintain the *Daphnia* heat stress transcriptional signal, tubes were flash frozen in liquid nitrogen for 2 seconds, and immediately stored at −80 °C until RNA extraction. The following system was used for sample identification: RNAtype_Temperature&TankID (e.g. sample “eRNA_20T1’’ refers to eRNA collected at 20 °C from tank 1).

### Extractions of eRNA and oRNA samples

Samples of eRNA and oRNA were extracted using the RNeasy Mini Kit (Qiagen), following the manufacturer’s protocol with the following modifications. eRNA samples were thawed on ice, gently vortexed for 5 seconds and then centrifuged for 3 minutes at 13,300 rpm to separate the liquid from filter. To increase the eRNA yield, liquid from two technical replicates was transferred into a 5 mL microcentrifuge tube containing 800 μL 70% EtOh and pipette mixed. This mixture was transferred to an RNeasy spin column (700 μL at a time) and centrifuged at 10,000 rpm for 15 seconds. We repeated this four times, until all remaining liquid from the 5 mL microcentrifuge tube was transferred to the spin column. For oRNA, *Daphnia* samples were thawed on ice and homogenized via sterile pestle and mortar mixer for 60 seconds. We then added 400 μL of 70% EtOh to the homogenized tissue. The resulting mixture was pipette mixed and transferred to an RNeasy spin column, where it was centrifuged at 10,000 rpm for 15 seconds. The remainder of the eRNA and oRNA extraction procedures followed the manufacturer’s instructions, with RNA being eluted in 40 μL of DNase and RNase free molecular grade water.

### DNA digestion

Immediately following RNA extraction, an 18 μL aliquot of both RNA types underwent two rounds of DNA digestion using the DNA-*free*™ DNA Removal Kit (Invitrogen) which provided rDNase. We completed the first round of DNA digestion following the manufacturer’s instructions, but skipped the DNase inaction step. To ensure no DNA carryover, we completed a second round of DNA digestion as follows: 2.7 μL of DNase I Buffer and 1 μL rDNase were added and pipette mixed into each sample and the plate was gently vortexed, centrifuged, and then incubated at 37 °C for 20 minutes. Following incubation, 2 μL of DNase Inactivation Reagent was added to each well. The plate was incubated for 2 minutes at room temperature, and centrifuged at 3700 rpm for 5 minutes to pellet the DNase Inactivation Reagent. The supernatant containing RNA was transferred to individual sterile microcentrifuge tubes and stored at −80 °C until library preparation.

### RNA-seq library preparation and sequencing

We prepared libraries for whole transcriptome shotgun sequencing (RNA-seq) using the Illumina Stranded Total RNA Prep with Ribo-Zero Plus kit (rRNA depletion) and IDT^®^ for Illumina^®^ RNA UD Indexes Set A, following the manufacturer’s instructions. Equal volumes of RNA (post 2 rounds of DNA digestion) from each technical replicate were pooled and used as input material (250 ng and 26.4-65.56 ng for oRNA and eRNA, respectively) to prepare one biological replicate library per tank. During library amplification, we used 12 and 17 PCR cycles for oRNA and eRNA, respectively, which followed the instructions outlined by the manufacturer for the respective RNA input amounts. Library quantification, quality control and equimolar pool sequencing of the 16 libraries was conducted at the McGill Genome Centre on one Illumina NovaSeq 6000 S4 lane using paired-end 100 bp reads.

### Contamination prevention

We submerged all experimental and sampling equipment in a 20% bleach solution for 15 minutes and rinsed thoroughly five times with distilled water prior to use. All eRNA samples were processed in a pre-PCR clean laboratory dedicated to low quality environmental and ancient DNA/RNA samples. Prior to entering the pre-PCR sterile laboratory, researchers entered a decontamination room to change into the dedicated clean lab coats and put on face masks, hairnets and lab clogs with shoe covers. The laboratory workbench was soaked in a 20% bleach solution for 10 minutes and RNase WiPER™ (iNtRON Biotechnology) and 20% bleach was used to thoroughly wipe pipettes, vortex mixers and centrifuges. Aerosol filter barrier pipette tips were used to prevent cross-contamination of samples. Filtration negative control samples and molecular negative control samples (consisting of reagents or DNA/RNA free molecular grade water) were processed at each step alongside eRNA samples, and were free of contaminating nucleic acids as identified by failed PCR amplification using universal COI primers known to amplify *Daphnia* DNA (Leray et al., 2013). Successful DNA digestion of each eRNA and oRNA sample was verified by failed PCR amplification of RNA post-DNA digestion x2 using universal COI primers known to amplify *Daphnia* DNA (Leray et al., 2013). RNA-seq libraries were prepared from filtration and molecular negative control samples and were free of contamination as indicated by quality control checks and failed amplification conducted at the McGill Genome Centre.

### Bioinformatic pipeline and statistical approaches

Raw FASTQ files underwent initial sequencing quality inspection using FastQC (Andrews, 2010). Low quality sequences and adapters were removed using Trimmomatic (Bolger et al., 2014) (ILLUMINACLIP: lluminaStrandedTotalRNA_adapter.fa:2:30:15 TRAILING:30 HEADCROP:1 MINLEN:90). All statistical analyses were conducted using R (R Core Team, 2021, R version 4.1.2). With default parameters, STAR (Dobin et al., 2013) was used to map all eRNA and oRNA reads that passed quality control and trimming to the *Daphnia pulex* reference genome (Z. Ye et al., 2017). FeatureCounts (Liao et al., 2014) was used on the BAM file output from STAR to quantify gene counts. Only those genes with a sum of ten or more counts in either 20 or 28 °C were considered detected and used in the subsequent differential expression (DE) analyses, for both eRNA and oRNA.

DE analysis was conducted with DESeq2 (Love et al., 2014), comparing all 20 °C samples versus all 28 °C samples, for both eRNA and oRNA samples. To determine genes as significantly differentially expressed, we used a false discovery rate (FDR) adjusted p-value < 0.05. A Pearson’s chi-squared test with Yates’ continuity correction was conducted to test for an association between the significantly differentially expressed genes identified in eRNA and oRNA. WEGO 2.0 (J. Ye et al., 2018) was used to classify GO terms (acquired from the reference genome) to all annotated *D. pulex* DEGs. Gene ontology enrichment analysis was conducted using topGO (Alexa & Rahnenfuhrer, 2022) by comparing GO terms of the relevant genes to the genomic background. A FDR corrected p-value < 0.05 indicated significant enrichment. A principal component analysis (PCA) of the regularized log (rlog)-normalized eRNA and oRNA gene expression profiles (300 most variant genes) at both optimal and heat stressed conditions was conducted using DESeq2’s plotPCA function (Love et al., 2014). Heatmaps showing the relative expression (Z-score calculated for each gene) of all significantly differentially expressed genes between 20 and 28 °C, in both *D. pulex* oRNA and eRNA were generated using pheatmap (Kolde, 2019).

We used the SqueezeMeta metatranscriptomics pipeline for the eRNA community-wide analysis (Tamames & Puente-Sánchez, 2019). Briefly, metatranscriptomes were assembled in sequential mode and contigs were assembled using Megahit (Li et al., 2015). Diamond was used to align all contigs to the GenBank nr and KEGG databases for taxonomic assignment and KEGG ID annotation, respectively (Buchfink et al., 2015; Kanehisa & Goto, 2000). We performed differential gene expression analyses as described above for *Daphnia*, after we mapped, using Kallisto (Bray et al., 2016), eRNA reads to the reference genomes and transcriptomes of species known a priori to persist in the communities (*A. falcatus, S. quadricauda*, and *P. subcapitata*) and highly abundant species identified by SqueezeMeta (*Brachionus plicatilis*, *Volvox cateri*, *Stylonychia lemnae*, *Chlamydomonas eustigma*) (Aeschlimann et al., 2014; Han et al., 2019; Hirooka et al., 2017; Nag Dasgupta et al., 2018; Prochnik et al., 2010; Schomaker & Dudycha, 2021; Suzuki et al., 2018). We used the SQMtools R package to further analyze the SqueezeMeta data and created stacked-bar plots to represent the most relatively abundant taxa (Puente-Sánchez et al., 2020). We conducted non-metric multidimensional scaling analysis of all KEGG IDs to visualize the community-wide functional profiles.

## Results

### Daphnia pulex *gene detection and expression profiles*

All *Daphnia* reads within environmental RNA (eRNA) samples are of extra-organismal origin as we pre-filtered tank water at 60 μm to remove *Daphnia*. Sequencing eRNA from tank water and organismal RNA (oRNA) from *Daphnia* tissue detected 3,919 (21%) and 17,244 (93%) genes, respectively, of the 18,449 genes that compose the *Daphnia pulex* reference genome (Z. Ye et al., 2017). All *Daphnia* genes detected in eRNA were also detected in oRNA. A principal component analysis of the regularized log normalized expression for the 300 most variable genes showed separation between experimental groups (Fig. 1). The samples were separated across the first principal component (PC1) by RNA type and across the second principal component (PC2) by temperature, clustering eRNA, oRNA, 20 °C and 28 °C within their respective groups. The only exception was 20 °C eRNA sample T1, which unexpectedly clustered closely with the 28 °C eRNA group. PC1 and PC2 accounted for 64% of explained variance of all principal components (Fig. S1).

**Fig. 1.**
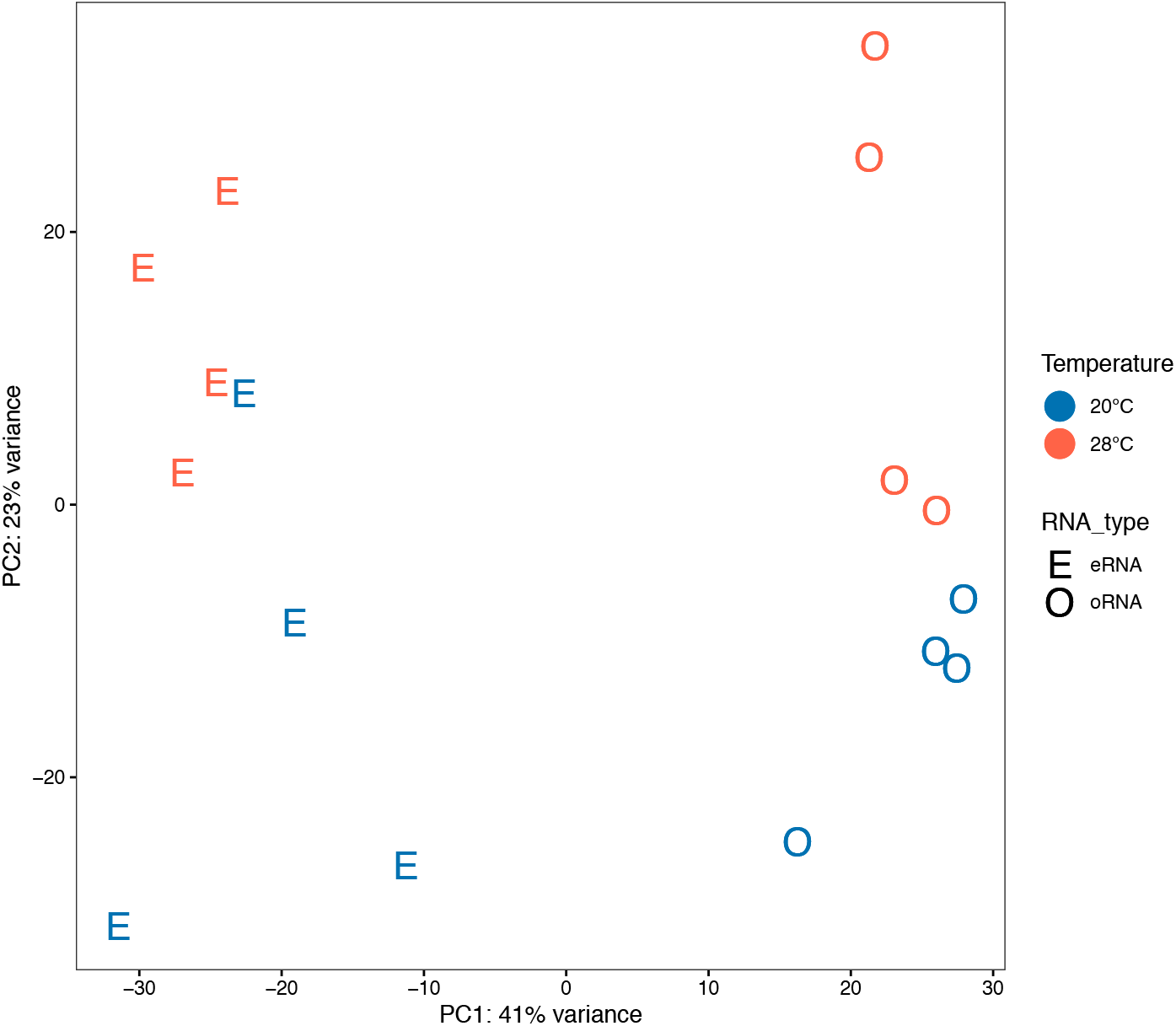
Principal component analysis (PCA) of the top 300 *Daphnia pulex* genes, as ranked by variance. eRNA refers to extra-organismal RNA released by *D. pulex* into the environment, whereas oRNA refers to organismal RNA extracted directly from *D. pulex* tissue. Prior to the PCA, genes were filtered to include only those that had a sum > 10 counts in either 20 °C or 28 °C. PCA revealed separation by RNA type and temperature across the PC1 and PC2, respectively (except eRNA_20T1 clusters more closely with 28 °C samples than with other 20 °C samples).

### Daphnia pulex *differential gene expression*

Comparison of gene expression profiles at 20 °C and 28 °C identified 32 significantly differentially expressed genes (DEG; false discovery rate adjusted p-value <0.05) from eRNA (4 upregulated and 28 downregulated) and 2351 DEGs from oRNA samples (1105 upregulated and 1246 downregulated). All eRNA significant DEGs were substantially up and down regulated (exceeding Log2 2 fold change), as well as 41 and 202 oRNA DEGs, respectively. The same directional change (up/down regulated) was observed in 31/32 eRNA DEGs as in oRNA (Table S1). Of all eRNA DEGs, 17 (2 upregulated and 15 downregulated) were also significantly differentially expressed in oRNA (Fig. S2). There was a significant association between DEGs identified in oRNA and DEGs identified in eRNA (χ2 = 18.26, df = 1, p-value = 1.92e-05). A heatmap comparing the relative expression (Z-score) of the 17 *D. pulex* DEGs common to eRNA and oRNA revealed four distinct clusters based on hierarchical clustering analyses, separating samples and genes by temperature and directional change, respectively (Fig. 2). Across both temperature conditions, eRNA and oRNA clustered together and were not separated into RNA types, indicating that eRNA and oRNA exhibit similar levels of relative gene expression for these DEGs. The only exception was eRNA 20 °C T1 sample which, similar to the PCA, unexpectedly clustered with the 28 °C samples.

**Fig. 2.**
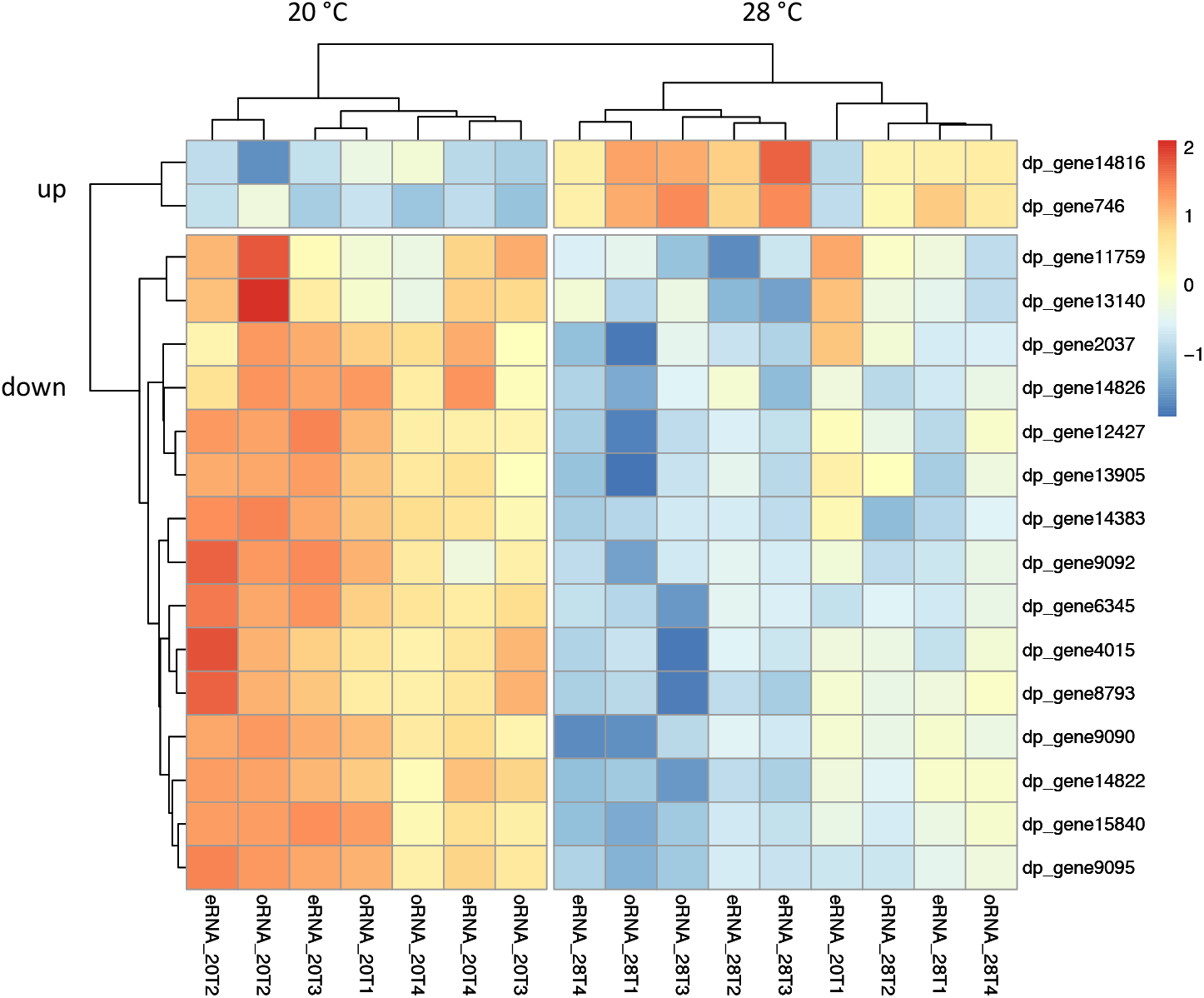
Heatmap showing the relative expression (Z-score calculated for each gene) of all commonly significantly differentially expressed (DE) *Daphnia pulex* genes (false discovery rate adjusted p-value <0.05) between 20 °C and 28 °C, in both environmental RNA (eRNA) and organismal RNA (oRNA). DE analysis was conducted only for those genes that had > 10 counts in either 20 °C or 28 °C samples. Hierarchical clustering analysis revealed four groups: 20 °C and 28 °C samples, and up and down regulated genes (except sample eRNA_20T1 clusters with 28 °C samples). Colors indicate levels of relative expression, with blue and red indicating low and high, respectively. DE statistics and functional description of the genes are provided in Table S1.

### Daphnia pulex *functional annotation*

Gene ontology (GO) enrichment analysis of eRNA detected genes identified 50 GO terms as enriched (FDR-corrected p-value <0.05), with 26, 6, and 18 belonging to Biological Processes (BP), Cellular Component (CC) and Molecular Function (MF) domains, respectively (Figs. 3 and S3). Of the 32 significant *D. pulex* DEGs identified from eRNA, 21 had GO annotations from the reference genome (Z. Ye et al., 2017) (Fig. S4-5). GO enrichment analysis of highly DE (exceeding Log2 2 fold change) eRNA genes, and DEGs common to both eRNA and oRNA, identified structural constituent of cuticle and chitin metabolic process, as enriched, respectively (FDR-corrected p-value <0.05; Fig. 3).

**Fig. 3.**
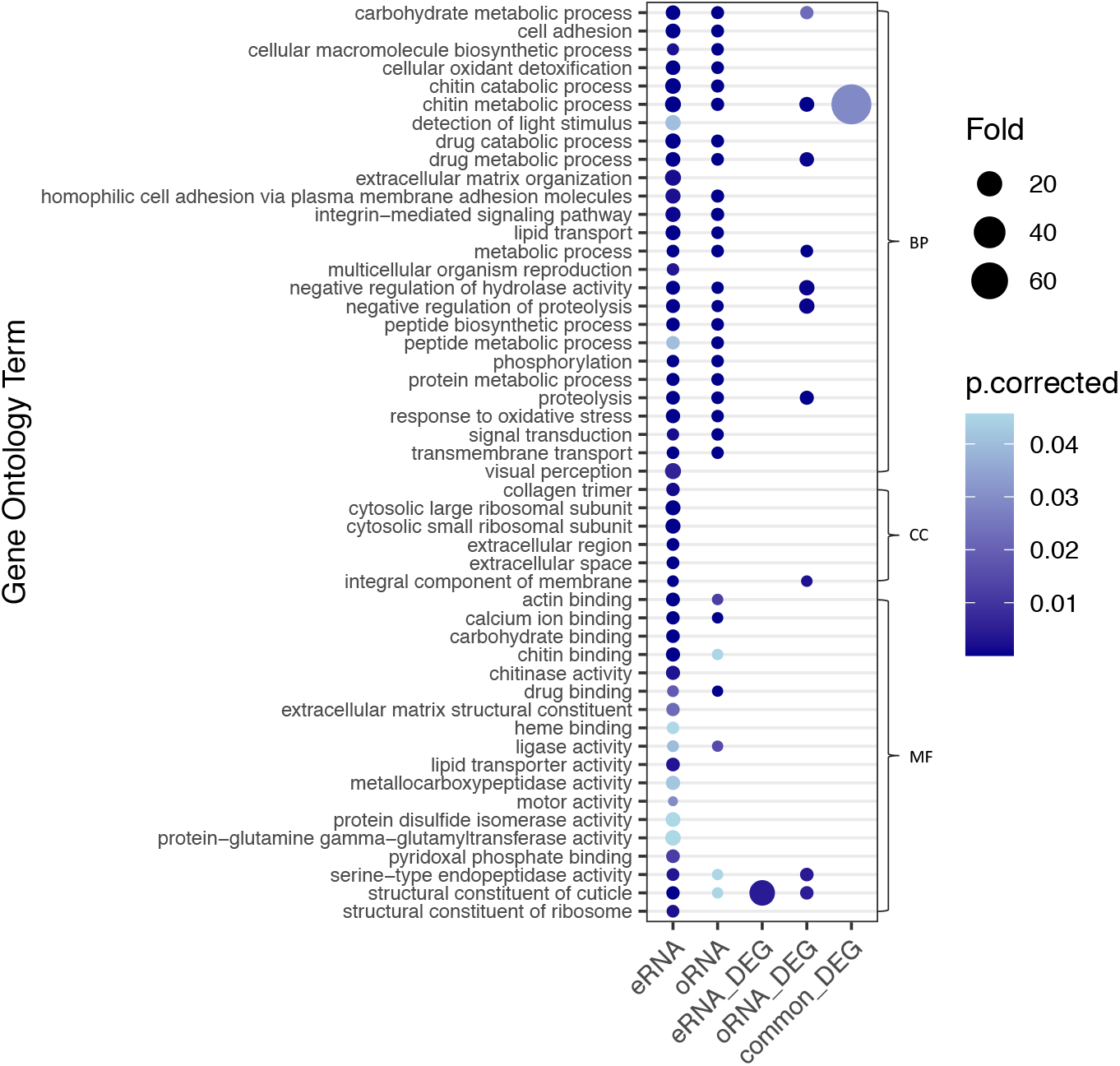
Bubble plot of the Gene Ontology (GO) enriched terms associated with *Daphnia pulex* eRNA and oRNA detected genes, and eRNA, oRNA and commonly identified differentially expressed genes (DEG). The eRNA genes originate from extra-organismal *Daphnia* RNA. We used a false discovery rate (FDR) corrected p-value <0.05 to identify GO terms as significantly enriched compared to the genomic background. The bubble color indicates the FDR corrected p-value for the weight test statistic, while the size is proportional to the fold change between expected and observed genes annotated with GO term. The biological processes, cellular components and molecular functions GO domains are represented by BP, CC, and MF, respectively. Only those GO terms enriched for eRNA genes are shown. Full list of GO terms enriched for oRNA genes is provided in Fig. S3.

GO enrichment analysis of oRNA detected genes identified 114 GO terms as enriched (FDR-corrected p-value <0.05), with 100, 1, and 13 belonging to BP, CC and MF domains, respectively (Figs. 3 and S3). Of the 2351 DEGs identified from oRNA, 1686 had GO annotations from the reference genome (Z. Ye et al., 2017) (Figs. S4-5). GO enrichment analysis of highly DE (exceeding Log2 2 fold change) oRNA genes identified 16 GO terms as enriched (FDR-corrected p-value <0.05), including structural constituent of cuticle, chitin metabolic process, proteolysis, negative regulation of hydrolase activity, negative regulation of proteolysis and response to stimulus (Fig S3).

### Community composition, differential gene expression and functional annotation

eRNA reads originated from a simple mock freshwater community that we constructed, consisting of *Daphnia pulex* as well as three phytoplankton species and opportunistic microorganisms that colonized the artificial lake water. All non-*Daphnia* eRNA reads likely originate from whole microorganisms that were captured on the filter. We used the SqueezeMeta pipeline (Tamames & Puente-Sánchez, 2019) to align eRNA reads to the GenBank nr database using its LCA algorithm, which identified 434 taxa. The community composition and relative abundances remained similar between 20 °C and 28 °C conditions (Fig. S6). Across eRNA samples, an average of 36% of all reads were aligned to eight species including four eukaryotes that were a priori known to persist in the communities and four highly abundant taxa identified by the SqueezeMeta analysis (Table S2). We detected 10,026 genes from these eight species, with the majority originating from *D. pulex, Scenedesmus quadricauda and Ankistrodesmus falcatus*, with 3919, 3180 and 1982 genes, respectively (Table S3). Differential gene expression analysis between 20 °C and 28 °C eRNA samples was performed for each of the eight species and identified a total of 121 significantly differentially expressed genes (FDR-adjusted p-value < 0.05). The highest numbers of DEGs were identified for *A. falcatus, D. pulex* and *S. quadricauda* with 44, 32 and 20 DEGs respectively.

Within all eRNA reads, 12,211 KEGG IDs were identified and a non-metric multidimensional scaling plot of these KEGG IDs revealed distinct functional profiles between 20 °C and 28 °C communities (Fig. S7). Of all KEGG IDs, 93 were identified as significantly differentially expressed (DE; false discovery rate-adjusted p-value <0.05) between 20 °C and 28 °C (Table S4). This included KEGG IDs corresponding to proteins and pathways involved in stress responses, such as the p53 signaling pathway, ubiquitin mediated proteolysis, metabolic pathway and cyclin-dependent kinase 4 (Table S4).

## Discussion

The use of environmental transcriptomics based on eRNA remained previously untested despite recent evidence demonstrating that extra-organismal eRNA persists in the environment long enough to be extracted (Kagzi et al., 2022; Littlefair et al., 2022; Marshall et al., 2021; Tsuri et al., 2021; Wood et al., 2020). Our findings revealed that environmental transcriptomics is sensitive in detecting transcriptional heat stress responses even without sampling source organisms directly. From eRNA released by *Daphnia pulex* into the tank water, we detected thousands of *D. pulex* genes and identified a subset of heat stress relevant genes to be differentially expressed between the two temperature conditions, with levels of relative expression similar to organismal RNA (oRNA). We also detected community wide changes in functional profiles. Our study demonstrates the ability of environmental transcriptomics to reveal gene expression responses of macroorganisms, and potentially complex biotic communities, following environmental changes.

### *Recovery of* Daphnia pulex *extra-organismal eRNA*

Several studies have recently demonstrated robust detection and persistence of extra-organismal eRNA in the environment (Kagzi et al., 2022; Marshall et al., 2021; Wood et al., 2020). We similarly were able to recover *Daphnia* eRNA in all samples, and had an average of 0.51% of all eRNA reads mapping to the *D. pulex* reference genome (Table S2). Our recovery of extra-organismal eRNA reads was similar to metagenomic studies of natural aquatic ecosystems that found between 0.27% and 1.25% of environmental DNA (eDNA) reads to be likely of extra-organismal origin (Cowart et al., 2018; Monchamp et al., 2022; Stat et al., 2017; Székely et al., 2021). We pre-processed our libraries with rRNA depletion, but could have potentially increased our recovery of *Daphnia* eRNA reads by targeting eukaryotic RNA through poly(A) enrichment. Nevertheless, our recovery of eRNA was particularly notable, given the general expectation that within environmental samples, potentially low-abundance eRNA is ‘competing’ to be sequenced with larger quantities of oRNA from whole microorganisms. This encouraging finding suggests that most of the eRNA captured is in a protected state (e.g., cellular, vesicular, or in stable molecular complexes) while free RNA forms, prone to degradation, are very short lived and potentially undetectable.

### Daphnia pulex *extra-organismal eRNA differential gene expression analysis*

Our broad environmental transcriptomics approach was able to detect 3,919 *D. pulex* genes from eRNA, building upon targeted RT-PCR of eRNA which detected 6 fish genes (Tsuri et al., 2021). As expected, we identified many more (2,351) differentially expressed genes (DEGs) under temperature stress from *Daphnia* oRNA than eRNA (32). This discrepancy in DEG identification can be easily attributed to large differences in the amount of RNA captured that is specific to *Daphnia* among other organisms in the community. This is reflected in the number of reads, 1 and 200 million, that mapped to the *D. pulex* reference genome using eRNA and oRNA, respectively (Table S2). Having less than 1% of eRNA reads matching *D. pulex* limited our power to detect DEGs, but is expected given that the libraries were generated from diverse environmental samples which included RNA transcripts from over 400 taxa, whereas the oRNA libraries were prepared from concentrated *D. pulex* tissue samples. On the other hand, we believe this diversity of eRNA within water samples could be beneficial as it opens up exciting possibilities of conducting community-wide surveys. For our proof-of-concept study, we performed rRNA depletion and deep sequencing (average of 158 million reads per eRNA sample, Table S2) in anticipation of such diverse eRNA samples to enable sufficient recovery of *D. pulex* eRNA for gene expression analysis. Despite the lower abundance of *D. pulex* reads from eRNA than oRNA, 97% of eRNA DEGs exhibited the same directional change (up/down regulated) as in oRNA (Table S1). We found a significant association between the DEGs identified in oRNA and those identified in eRNA, and similar levels of relative expression in commonly shared DEGs (Fig 2). These results demonstrate that similar to conventional oRNA tissue samples, our environmental transcriptomics approach yielded adequate sensitivity in detecting changes in gene expression of a source organism in response to heat stress.

Changes in gene expression are often observed before phenotypic and demographic effects occur and may therefore act as an early indication of ecological stress (Fedorenkova et al., 2010; Jovic et al., 2017; Snell et al., 2003). We found that eRNA and oRNA recovered similar *D. pulex* chronic heat stress responses which consisted of extensive downregulation of genes associated with metabolic and cellular processes, membrane, binding, extra-cellular region, collagen and cuticle structure (Figs. 3 and S4). These results are consistent with studies that also found chronically heat stressed *Daphnia* to exhibit widespread downregulation of genes with similar functions (Becker et al., 2018; Yampolsky et al., 2014). This widespread downregulation of metabolic genes has been hypothesized to be a molecular compensatory mechanism for *Daphnia* at near-lethal temperatures to sacrifice their long-term fitness for immediate survival (Yampolsky et al., 2014).

After seven days of heat exposure, only few (4/62) heat shock genes were slightly upregulated (log_2_ fold change =1.1-1.3; Table S5) in *Daphnia* oRNA. This aligns with findings suggesting heat shock genes to be less important for the chronic (days) *Daphnia* heat stress response, but central for mitigating cell damage following acute (hours) heat exposure (Becker et al., 2018; Yampolsky et al., 2014). Multiple tropical marine fish species similarly do not differentially express heat shock genes during chronic heat stress (Veilleux et al., 2015), suggesting that heat shock genes might not be optimal for assessments of communities under chronic heat stress. Furthermore, we detected chitin related gene ontology (GO) terms to be commonly enriched in both eRNA and oRNA DEGs (Fig. 3), corroborating *Daphnia* studies that linked chitin genes to environmental stressors (Becker et al., 2018; Chain et al., 2019; Connon et al., 2008; Shaw et al., 2007). Hydrolytic and proteolytic related GO terms were enriched in eRNA (Fig. 3), and are commonly involved in *Daphnia* and copepod heat stress response by stabilizing and cleaning up misfolded proteins (Becker et al., 2018; Kelly et al., 2017). It is promising that using eRNA, we were able to detect the widespread downregulation of genes associated with chronic heat stress with similar levels of relative expression as oRNA. We also found that many GO terms were commonly shared in both RNA types, and that stress related terms were enriched in eRNA. Collectively, our results indicate that environmental transcriptomics based on eRNA is sensitive in detecting heat stress responses of source organisms through changes in gene expression and functional GO terms. Future empirical work is necessary to determine how our results extend to more complex natural ecosystems.

### Community composition and function

We found the community composition to remain largely unchanged for 20 °C and 28 °C conditions, after mapping all eRNA reads to GenBank nr (Fig. S6). Since we pre-filtered the tank water at 60 μm to remove *Daphnia*, all *non-Daphnia* RNA recovered in our eRNA samples likely originates from whole microorganisms that passed the filter. Within our eRNA samples, gene expression analysis of eight species detected 10,026 genes with 121 of these being significantly differentially expressed (DE) between 20 °C and 28 °C (Table S3). Although community compositions remained similar, analysis of KEGG orthologs revealed distinct functional profiles for communities in the two temperature treatments (Fig. S7). We found stress-related KEGG orthologs to be DE, including proteins involved in binding, kinase and oxidoreductase (Table S4). Our community wide results reflect similar patterns of differential gene expression and shifts in functional terms observed in conventional aquatic metatranscriptomic studies based on bulk microorganism samples (Aylward et al., 2015; Frias-Lopez et al., 2008; Gilbert et al., 2008; Marchetti et al., 2012; Moniruzzaman et al., 2017; Poretsky et al., 2005; Salazar et al., 2019; Vorobev et al., 2020). For example, a freshwater metatranscriptome likely under heat stress exhibited similar stress responses which included oxidoreductase, binding and kinase proteins (Trench-Fiol & Fink, 2020). We build upon this rich metatranscriptomics literature by demonstrating that in addition to yielding microorganism insights, environmental transcriptomics can also non-invasively detect the gene expression response of macroorganisms to environmental stress.

### Recommendations and future directions

Interest in non-lethal transcriptomics is growing, as such an approach increases animal welfare, allows for repeated survey of individuals, and linking gene expression to fitness following exposure to stress (Czypionka et al., 2015; Jeffries et al., 2021; Veldhoen et al., 2014). However, conventional non-lethal methods are species specific and involve animal handling and sampling for blood or tissue, which may still result in organism stress and reduced survival upon release into the environment (Martins et al., 2018; Portz et al., 2006; Young et al., 2019). Inferring the physiological state of organisms from non-invasive eRNA samples would circumvent these stressful sampling procedures and represent a substantial improvement for animal welfare. We demonstrated that similar to the conventional oRNA approach, environmental transcriptomics based on eRNA can non-invasively detect heat stress responses of progenitor organisms.

An exciting prospect of environmental transcriptomics is the potential to monitor the ecological health of complex communities, including multiple macroorganisms. Biomonitoring traditionally relies upon identifying bioindicator taxa that are correlated with environmental conditions, but transcriptomic surveys can provide valuable early warnings since gene expression changes are likely to occur before compositional shifts could be detected (Veilleux et al., 2021). Conventional metatranscriptomics has demonstrated that gene expression profiling of microorganisms reflects environmental conditions (Marchetti et al., 2012; Moran et al., 2013; Salazar et al., 2019; Shi et al., 2011). Environmental transcriptomics extends beyond microorganisms and has the potential to provide functional information of species across the trophic chain as eRNA includes extra-organismal RNA from a diversity of macroorganisms. However, further empirical work is required before such community-wide surveys should be applied to biomonitoring efforts; population-level assessments that couple eRNA with targeted approaches (e.g. qPCR, ddPCR) and stress biomarkers are more realistically applicable in the short-term. Nevertheless, gaining insights into the ecological health of populations and communities would enable conservation initiatives to be proactive in their efforts and respond before local extinction events occur. Environmental transcriptomics is in its infancy and refinement through mesocosm experiments and natural environment studies must first be conducted before this approach can be applied further.

We anticipate multiple challenges for environmental transcriptomic biomonitoring in natural environments, such as achieving adequate sequencing depth of eRNA in a cost-efficient manner (Table 1). However, sequencing technologies are rapidly advancing and the costs continue to decrease (Wetterstrand, 2021). The continuation of such technological advancement will enable future studies to generate more reads less expensively, thereby increasing the sensitivity of environmental transcriptomics to detect extra-organismal eRNA transcripts. Additionally, the continued development of low input mRNA-seq library preparation kits will enable researchers to prepare libraries for environmental transcriptomics with minimal template eRNA. Molecular technological advancements that enhance extraction efficiency, enrich eukaryotic RNA and remove ribosomal RNA from diverse samples may increase the yield of coding eRNA and thus the power of environmental transcriptomics.

**Table 1.** Problems associated with using environmental transcriptomics based on eRNA for biomonitoring and potential experimental (E), molecular (M), sequencing (S) and bioinformatic (B) solutions.

| Problems | Description of problems | Possible solutions | References |
| --- | --- | --- | --- |
| Poor understanding of the ecology of eRNA | Little is known about the origin, state, persistence and degradation of eRNA in the environment. | E: Perform experiments that address fundamental questions of the ecology of eRNA (i.e. What are major sources of eRNA? What state is eRNA in? What factors influence eRNA persistence? How do abiotic factors influence eRNA decay rates?) | (Cristescu, 2019) |
| No understanding of environmental transcriptomics in natural ecosystems | No information available about how our results using environmental transcriptomics will translate to natural ecosystems | E: Perform mesocosm experiments using more complex, multi-trophic communities<br>E: Perform studies using ecosystem wide manipulations<br>E: Perform studies in natural ecosystems |  |
| Rapid degradation of eRNA | eRNA is thought to degrade too rapidly to extract and quantify, but studies indicate that eRNA is detectable for up to 57 hours post organism release | E: Filter water as soon as possible (ideally < 4 hours)<br>E: Freeze samples at -80 °C directly after filtration. Alternatively could potentially use preservative solution if freezing immediately is not possible.<br>M: Keep samples on ice during extraction<br>M: Minimize free-thaw cycles | (T. S. Jo et al., 2022; Kagzi et al., 2022; Marshall et al., 2021; Wood et al., 2020) |
| Low concentration of eRNA | eRNA samples have much lower RNA concentrations than samples collected from an organisms tissue (i.e. 1 ng/μL vs 200 ng/μL) | E: Increase RNA yield by filtering large volumes of water (> 500 mL);<br>EM: Collect additional technical replicates and pool them during the extraction step;<br>M: Use RNA-seq library prep kits designed for low input starting material |  |
| Low proportion of mRNA | mRNAs are of interest for gene expression analyses as they encode for proteins, but they account for < 5% of total RNA. Non-coding ribosomal and small RNAs account for > 85% | M: Perform rRNA depletion to remove rRNA. However, many commercial kits are optimized to remove human, mouse, rat and bacteria rRNA, and may not deplete rRNA from all species;<br>M: Perform poly(A) enrichment to retain only eukaryotic mRNA. This will eliminate prokaryotic RNA, ribosomal and small RNAs, which make up the majority of total RNA<br>B: If rRNA persists, use it to perform taxonomic identification analysis | (He et al., 2010; Hempel et al., 2022; S. Zhao et al., 2018; W. Zhao et al., 2014) |
| Low eRNA recovery | eRNA samples are composed of extra-organismal RNA from macro-organisms and whole microorganism. Thus, eRNA reads will be in the minority (estimated <1%), resulting in low power of bioinformatic analyses to detect eRNA based changes in gene expression | E: Pre-filter water with fine mesh to size-selectively remove organisms;<br>E: Use large filter pore size to reduce capture of microorganisms (i.e. > 0.2 μM);<br>E: Sample eRNA from ecosystems that are densely populated with target species<br>M: Perform rRNA depletion on total RNA samples;<br>M: Perform poly(A) enrichment to eliminate prokaryotic RNA;<br>S: Perform deep sequencing to increase eRNA reads (i.e. > 150 million reads per sample) |  |
| Low eRNA read depth | eRNA samples are extremely diverse, resulting in reads from taxa across the trophic chain. Transcript abundance in a sample will influence read depth, and if many micro-organisms are present, eukaryotic genes will have low read depth. Current molecular methods for enrichment and depletion are | M: Create low input poly(A) enrichment methods. Current enrichment methods are designed for organismal RNA from tissue samples and require high concentration of input RNA;<br>M: Increase eukaryotic gene read depth through poly(A) enrichment;<br>MB: Create rRNA depletion methods that remove rRNA from all taxa as current depletion methods are optimized to remove single taxa and bacterial rRNA. If unable to remove all rRNA from eRNA samples, researchers could use rRNA reads for taxonomic information;<br>M: Perform rRNA depletion; | (Yates et al., 2021) |
|  | optimized for organismal, not diverse environmental samples. | M: Amplify biomarker genes via targeted (qPCR, ddPCR) approaches to analyze specific biomarker genes |  |
| Difficulty characterizing read function due to lack of eukaryotic reference genomes | Reference genomes are used to functionally and taxonomically characterize reads. eRNA samples will capture eRNA from eukaryotes across the trophic chain, but genomes are available for only 0.4% of eukaryotic species. | ES: Sequence all eukaryotic species (e.g. Earth BioGenome Project)<br>B: *De novo* assembly of reads into longer scaffolds, which enables the identification of homologs. | (Lewin et al., 2018; Shakya et al., 2019) |
| Lack of baseline in natural ecosystems to compare against | Differential gene expression analyses are conducted by comparing treatment samples (i.e. stressed community) against a control 'baseline' (i.e. health community). In natural ecosystems it is difficult or often not possible to obtain a baseline control as many stressors act simultaneously on a system | E: Conduct time series eRNA sampling of the same environment and compare time points;<br>E: Compare environments with similar abiotic conditions. If healthy, one could serve as a baseline to compare against;<br>EM: Couple eRNA with targeted PCR approaches (qRT-PCR, ddPCR) and biomarker genes to infer physiological states of specific species;<br>EB: Reconstruct mock communities under neutral conditions as a baseline;<br>B: The cellular stress response is triggered by general stress and as such genes and pathways involved could be broadly indicative of community stress;<br>B: Use housekeeping genes with consistent expression levels as baselines to compare against;<br>B: Supervised machine learning models could be trained with existing metatranscriptomic, transcriptomic and metagenomic data that have identified functional terms, metabolic pathways and taxa associated with stressors. Environmental transcriptomic data could be used as input and the model would predict the stressors involved and health of the community | (Cordier et al., 2019; Fulda et al., 2010; Mallick et al., 2021; Yates et al., 2021; Zhang et al., 2021) |
| False detection of differential transcript expression due to changes in population densities | Between environments, species across the trophic chain exhibit differences in abundances. RNA transcript abundance varies with DNA abundance. Thus, changes in RNA transcript abundance, independent of differential expression, could be recovered. | EB: Collect and sequence eDNA in parallel. eDNA can act as a species abundance estimator, independent of transcript abundance, that we can normalize eRNA reads to. Allows us to track whether the proportion of eRNA reads follows species abundance, or is up or down regulated;<br>B: 'Taxon-specific scaling' accounts for variation in abundances by separating and then normalizing metatranscriptomic data into organism-specific bins. A differential expression analysis could then be performed after the normalized matrices are recombined into community wide data | (Klingenberg & Meinicke, 2017; Salazar et al., 2019; Yates et al., 2021; Zhang et al., 2021) |

While sequencing and molecular advancements will resolve challenges associated with eRNA depth, perhaps the largest remaining challenge for environmental transcriptomics to overcome is the lack of a baseline to compare against. Differential gene expression analyses are conducted by comparing a treatment against a control ‘baseline’ to identify genes involved with the stress response. However, in natural ecosystems, it is difficult to obtain a control baseline due to population demographics and multiple stressors acting simultaneously on a system. This is similarly shared by metatranscriptomics, and multiple statistical and mathematical modelling solutions have been developed to overcome this problem (Klingenberg & Meinicke, 2017; Mallick et al., 2021; Zhang et al., 2021).

By providing functional insights in a non-invasive manner, eRNA based approaches could push beyond the limitations of eDNA species detection and conventional tissue-based transcriptomic surveys. We demonstrated that environmental transcriptomics can detect thousands of genes, and that a subset of functionally relevant genes can be identified as differentially expressed under heat stress, with levels of relative expression similar to the conventional oRNA approach. Within all eRNA reads, we were able to reveal changes in functional profiles for the community in response to heat stress. Collectively, these findings demonstrate the ability of eRNA to detect changes in gene expression following environmental stress and the potential of environmental transcriptomics to provide functional information across the trophic chain. Future empirical work is necessary to determine if this approach can be extended to natural ecosystems.

## Data accessibility

Sequences for all RNA-seq libraries have been uploaded to the NCBI SRA database and will be made public upon acceptance of this manuscript. Reviewers can preview the uploaded files: dataview.ncbi.nlm.nih.gov/object/PRJNA830892?reviewer=mcvpem4l5cupljoaj9jn53kg1e

## Author contributions

RMH and MEC designed the study. RMH and MCY conducted the experiment. RMH conducted all molecular work. RMH and FJJC performed the bioinformatic analysis and produced the figures. RMH wrote the first draft of the manuscript and all authors contributed to editing the manuscript.

## Acknowledgements

We thank Elif Irmak Bektas, Cathy Shen and Gabrielle Pellegrin for assisting with animal husbandry and sample collection. We are grateful to Lars Iversen and Caren Helbing for providing comments on an early version of this manuscript and Takahiro Maruki for input regarding the bioinformatic analysis. This study was supported by the Natural Sciences and Engineering Research Council (NSERC) of Canada through the Canada Research Chair program and a Discovery Grant to Melania Cristescu.

