## Supplementary Information for "Environmental transcriptomics under heat stress: Can environmental RNA reveal changes in gene expression of aquatic organisms?"

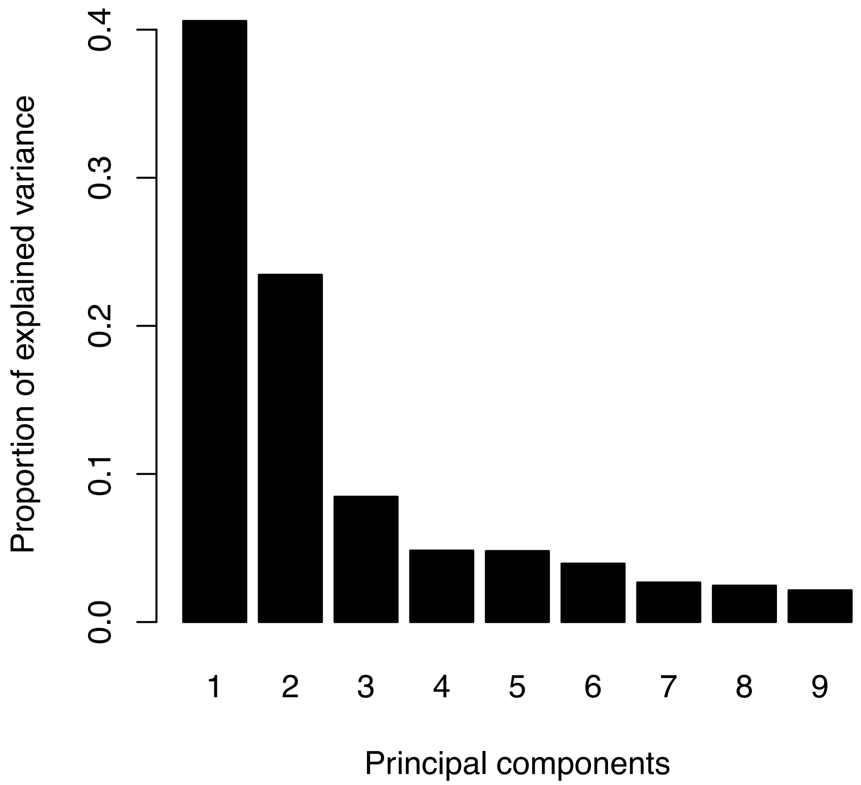


**Fig. S1.** Scree plot showing the proportion of variance captured by each of the top 9 principal components of Fig 1. The first and second principal components separate RNA type and temperature conditions, respectively.

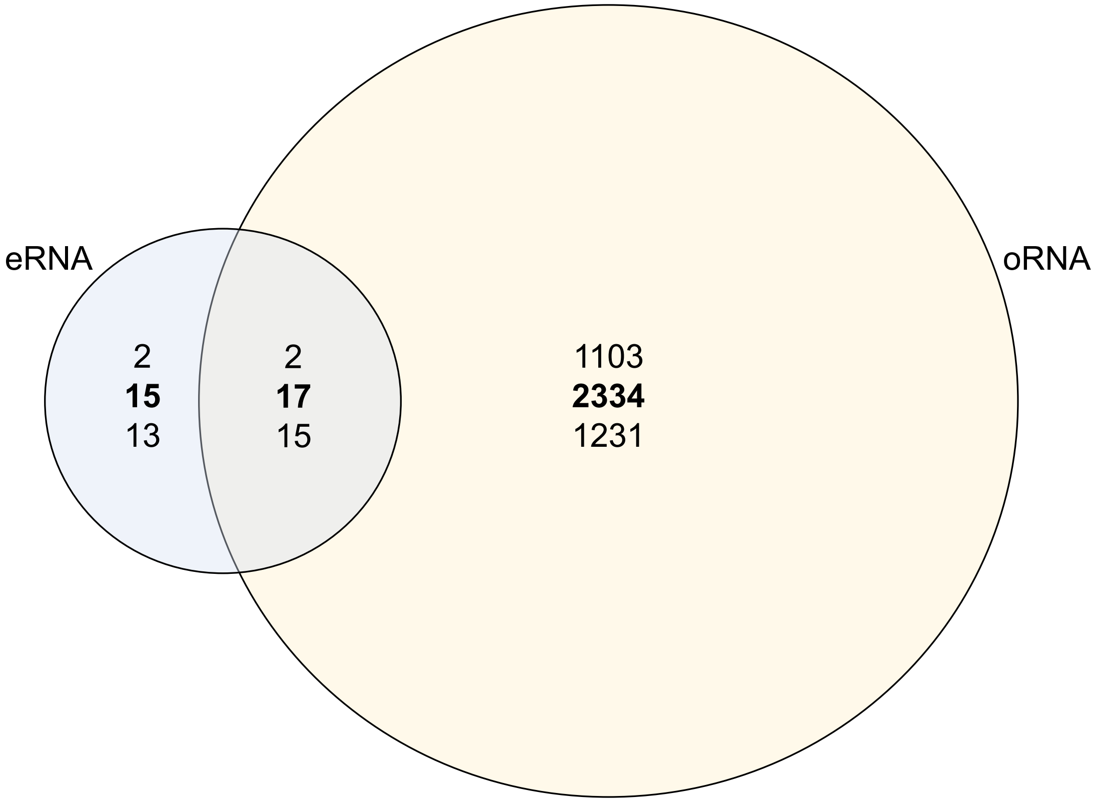


**Fig. S2.** Venn diagram showing the significantly differentially expressed *Daphnia pulex* genes (DEG; FDR-adjusted p-value $<0.05$) between 20°C and 28°C, in both eRNA (blue) and oRNA (yellow) samples. Bold number represents the sum of DEG, with up and down-regulated genes shown in brackets above and below, respectively. The genes within eRNA samples originated from extra-organismal RNA.

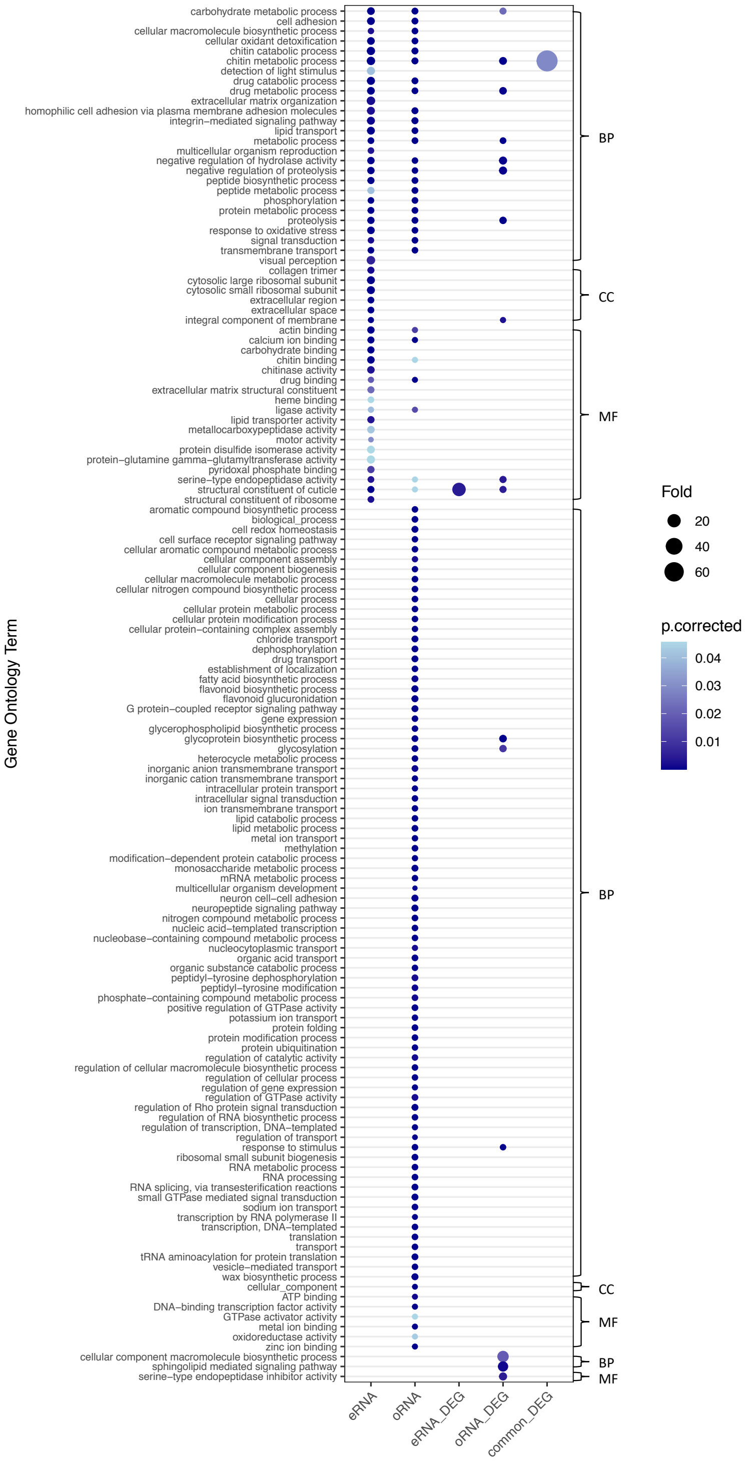


**Fig. S3** Bubble plot of the Gene Ontology (GO) enriched terms associated with *Daphnia pulex* eRNA and oRNA detected genes, and eRNA, oRNA and commonly identified differentially expressed genes. We used a false discovery rate (FDR) corrected p-value <0.05 in topGO to identify GO terms as significantly enriched compared to the genomic background.

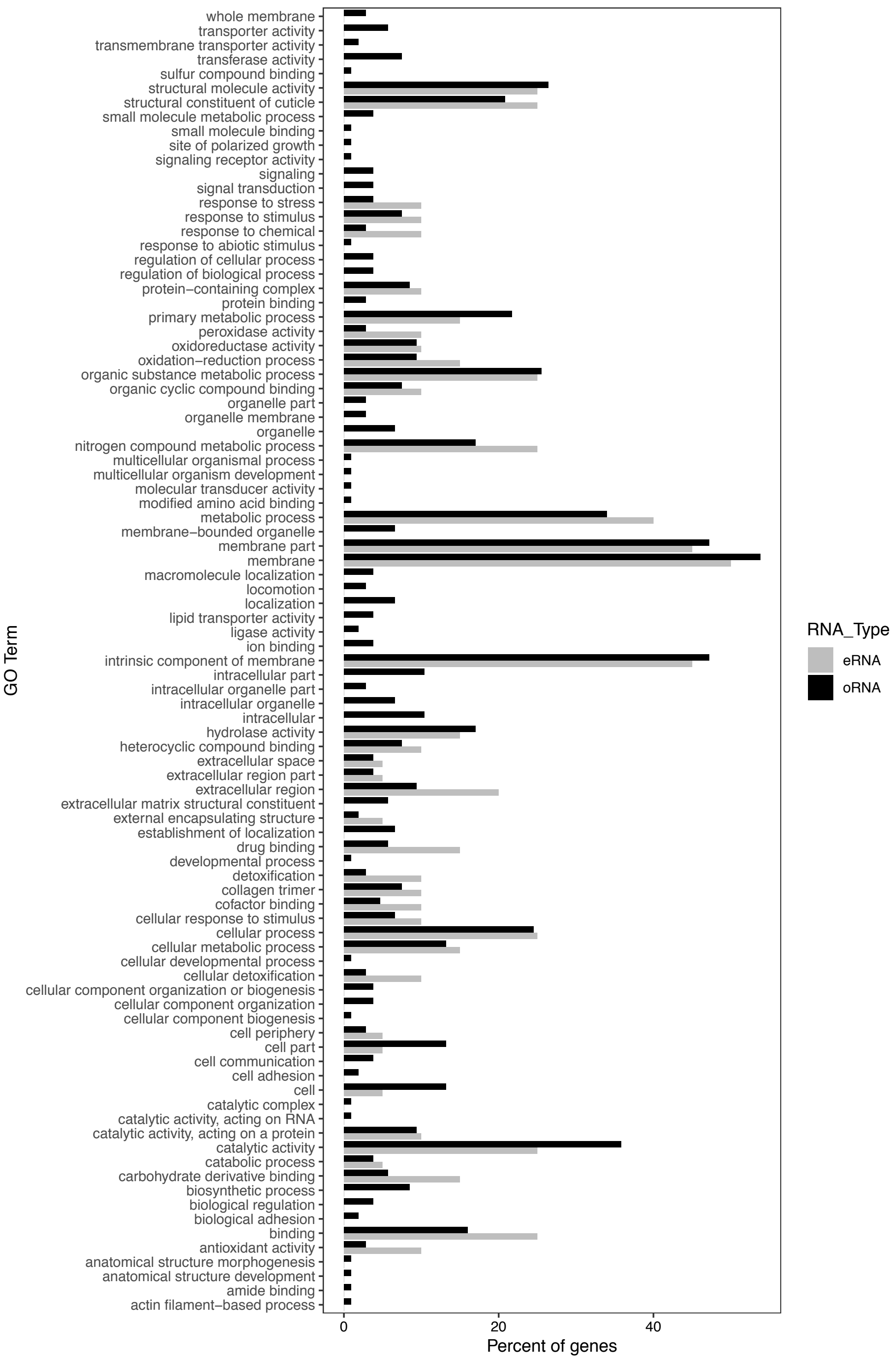


**Fig. S4.** The percent of eRNA (grey) and oRNA (black) *Daphnia pulex* significantly (FDR-adjusted p-value <0.05) and highly (log 2 fold change < -2) downregulated genes that were annotated with Gene Ontology (GO) terms. Note that the number of downregulated eRNA and oRNA genes annotated with GO terms varied and were 20 and 106, respectively.

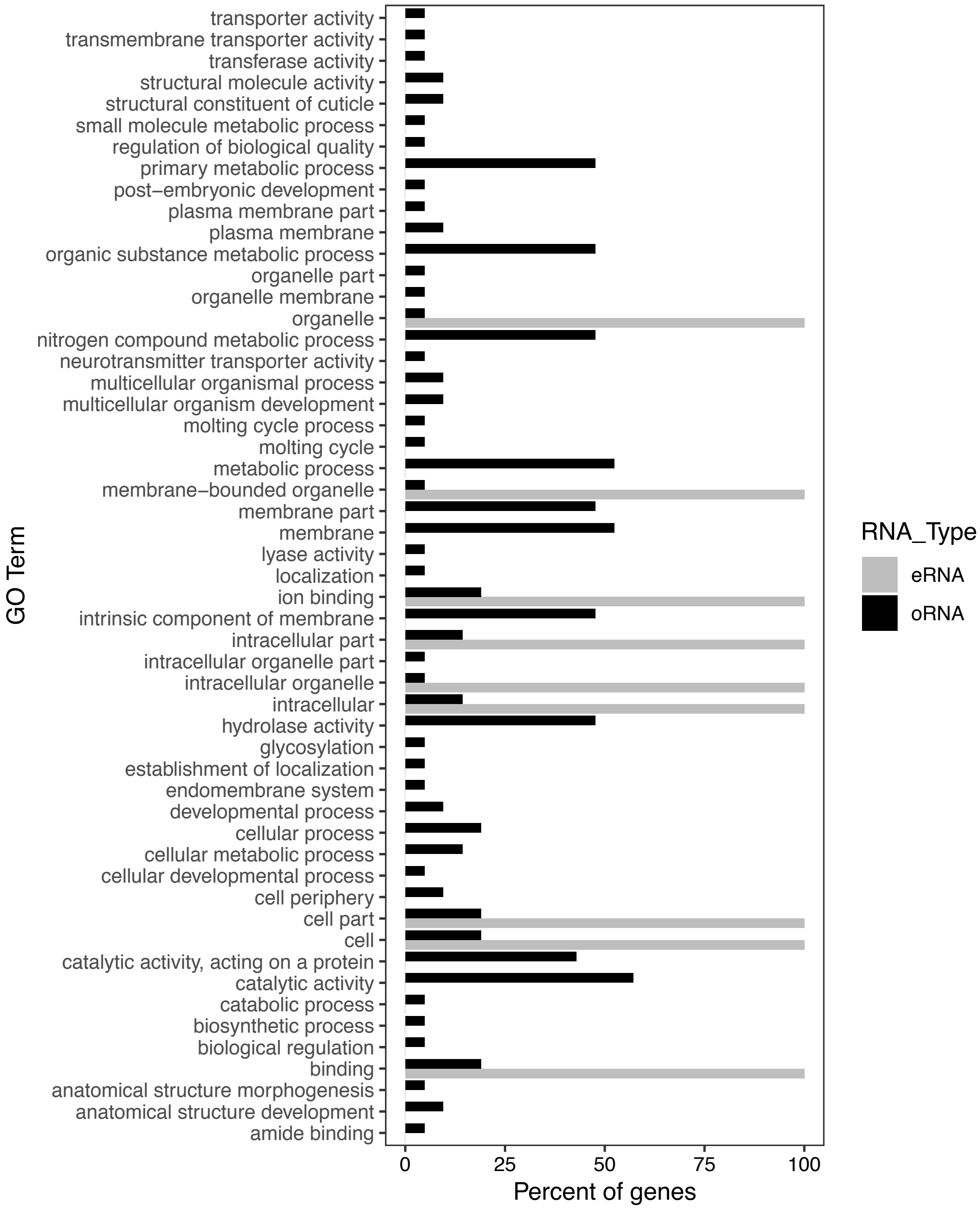


**Fig. S5.** The percent of eRNA (grey) and oRNA (black) *Daphnia pulex* significantly (FDR-adjusted p-value <0.05) and highly (log 2 fold change > 2) upregulated genes that were annotated with Gene Ontology (GO) terms. Note that the number of upregulated eRNA and oRNA genes annotated with GO terms varied and were 1 and 21, respectively.

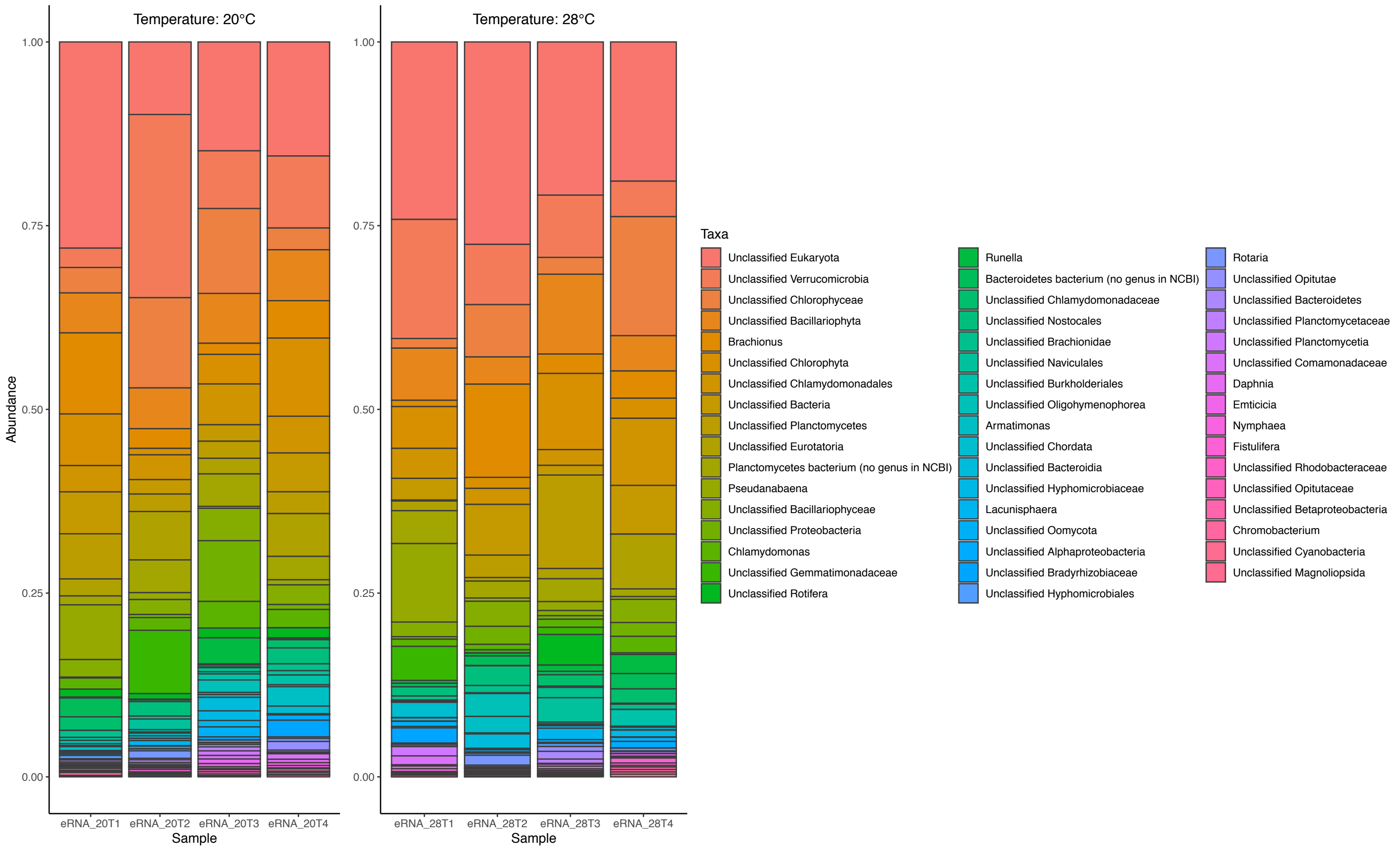


**Fig. S6.** Stacked bar-plot of the relative abundance for the 50 most abundant taxa in all eRNA samples. eRNA reads were aligned to the GenBank nr database using SqueezeMeta's last common ancestor (LCA) algorithm.

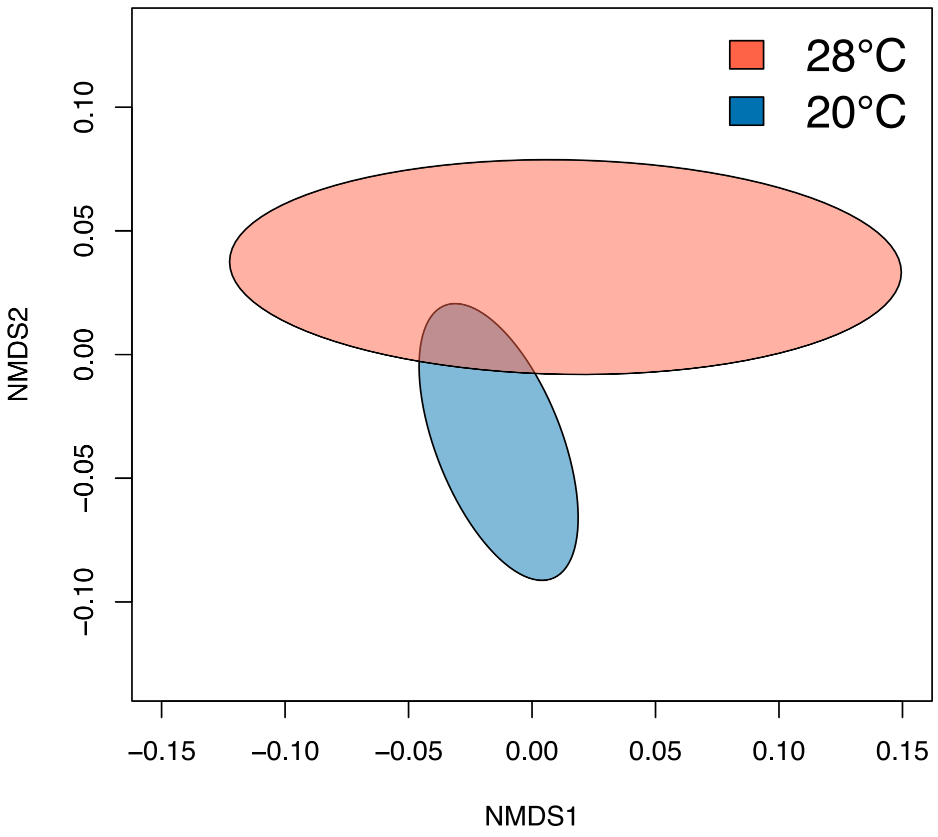


**Fig. S7.** Community wide functional profiles for the 20°C and 28°C eRNA samples represented by non-metric multidimensional scaling (NMDS) of functional KEGG orthologs. eRNA reads were aligned against the KEGG database using SqueezeMeta's classical best hit approach. KEGG orthologs differentially expressed between 20°C and 28°C eRNA samples are provided in Table S4.

**Table S1.** Statistics for all environmental RNA (eRNA) identified significantly expressed *Daphnia pulex* genes (false-discovery rate adjusted p-value <0.05). pvalue refers to the false-discovery rate adjusted p-value. LFC refers log2 fold change and positive and negative signs indicate up or down regulation, respectively. 31/32 genes show same direction of differential expression in both eRNA and oRNA, with the one exception being not significantly differentially expressed in oRNA. Functional descriptions were obtained from the *Daphnia pulex* reference genome (Ye et al. 2017).

| **Gene** | **eRNA LFC** | **eRNA pvalue** | **oRNA LFC** | **oRNA pvalue** | **Functional description** |
|---|---|---|---|---|---|
| dp_gene14816 | 7.04 | 0.0005 | 1.04 | 0.0041 | hypothetical protein DAPPUDRAFT_333097 |
| dp_gene2037 | -5.42 | 0.0042 | -2.35 | 0.0122 | DNA-directed RNA polymerase II subunit RPB1- |
| dp_gene15840 | -3.41 | 0.0042 | -2.08 | 0.0005 | Chitin deacetylase 1 precursor; extracellular region;integral component of membrane;chitin deacetylase activity;chitinase activity;chitin binding;carbohydrate metabolic process;chitin catabolic process |
| dp_gene9095 | -6.62 | 0.0042 | -2.08 | 0.0000 | Collagen iv alpha 1 chain;collagen trimer |
| dp_gene14383 | -5.90 | 0.0042 | -3.82 | 0.0000 | Flocculation FLO11; integral component of membrane |
| dp_gene13905 | -3.29 | 0.0042 | -1.58 | 0.0472 | Msr- isoform B; integral component of membrane |
| dp_gene4015 | -6.53 | 0.0050 | -1.93 | 0.0066 | Cuticular 50Cb; structural constituent of cuticle |
| dp_gene10667 | -5.21 | 0.0141 | -1.32 | 0.2518 | Cuticular 49Ag; extracellular space;structural constituent of cuticle |
| dp_gene746 | 5.84 | 0.0167 | 1.59 | 0.0000 | hypothetical protein DAPPUDRAFT_300235 |
| dp_gene12427 | -5.57 | 0.0167 | -2.34 | 0.0230 | hypothetical protein DAPPUDRAFT_220312 |
| dp_gene8793 | -4.61 | 0.0172 | -1.75 | 0.0096 | hypothetical protein DAPPUDRAFT_239781;integral component of membrane;structural constituent of cuticle |
| dp_gene4009 | -5.09 | 0.0172 | -1.50 | 0.2016 | Cuticle;integral component of membrane;structural constituent of cuticle |
| dp_gene13317 | -6.86 | 0.0172 | -1.73 | 0.1412 | Cuticle;integral component of membrane;structural constituent of cuticle |
| dp_gene8695 | -6.49 | 0.0210 | -2.37 | 0.0580 | hypothetical protein DAPPUDRAFT_315291;integral component of membrane |
| dp_gene7516 | -4.85 | 0.0226 | -1.80 | 0.1628 | hypothetical protein DAPPUDRAFT_101532 |
| dp_gene7027 | -5.18 | 0.0283 | -1.04 | 0.4070 | hypothetical protein DAPPUDRAFT_311888 |
| dp_gene13140 | -3.18 | 0.0283 | -1.99 | 0.0158 | hypothetical protein DAPPUDRAFT_240525;oxidation-reduction process |
| dp_gene12640 | 2.63 | 0.0283 | 0.31 | 0.5589 | hypothetical protein DAPPUDRAFT_307954 |
| dp_gene11759 | -2.49 | 0.0283 | -1.44 | 0.0380 | Trypsin serine protease;serine-type endopeptidase activity;proteolysis |
| dp_gene9092 | -5.92 | 0.0283 | -3.18 | 0.0000 | Collagen iv alpha 1 chain;collagen trimer |
| dp_gene14826 | -3.55 | 0.0319 | -2.10 | 0.0001 | chondroitin proteoglycan-2-like;extracellular region;chitin binding;chitin metabolic process |
| dp_gene12431 | -5.94 | 0.0388 | -2.09 | 0.1437 | hypothetical protein;integral component of membrane |
| dp_gene14822 | -4.47 | 0.0388 | -1.63 | 0.0020 | probable chitinase 3;extracellular region;chitin binding;chitin metabolic process |
| dp_gene4575 | -5.15 | 0.0409 | -1.56 | 0.1006 | Chorion peroxidase precursor;chorion;peroxidase activity;heme binding;response to oxidative stress;oxidation-reduction process;cellular oxidant detoxification |
| dp_gene21334 | -5.58 | 0.0419 | -2.34 | 0.0680 | hypothetical protein DAPPUDRAFT_101535 |
| dp_gene9090 | -3.64 | 0.0425 | -2.55 | 0.0009 | Collagen alpha-6(VI) chain |

| | | | | | |
|---|---|---|---|---|---|
| dp_gene2763 | -4.95 | 0.0425 | -1.33 | 0.3201 | Chorion peroxidase precursor;membrane;peroxidase activity;heme binding;response to oxidative stress;oxidation-reduction process;cellular oxidant detoxification |
| dp_gene6345 | -5.98 | 0.0425 | -2.57 | 0.0000 | Collagen iv alpha 1 chain |
| dp_gene1099 | 2.82 | 0.0435 | -0.54 | 0.1063 | LIM domain-binding;nucleus;zinc ion binding |
| dp_gene6653 | -3.96 | 0.0465 | -1.56 | 0.2076 | Spz1;integral component of membrane |
| dp_gene5484 | -3.28 | 0.0489 | -0.41 | 0.7236 | Gly d 3;serine-type endopeptidase activity;proteolysis |
| dp_gene7595 | -4.97 | 0.0489 | -1.91 | 0.1921 | hypothetical protein DAPPUDRAFT_316713 |

**Table S2.** Summary of alignment statistics for the total number of reads that mapped to the reference genomes of *Daphnia pulex, Ankistrodesmus falcatus, Raphidocelis subcapitata* and *Scenedesmus quadricauda* known a priori to persist in our tanks as well as four highly abundant taxa as identified by Diamond alignment to the NCBI nr database. Sequencing was conducted on one Illumina NovaSeq 6000 S4 lane with 100bp paired-end reads

| Sample ID | Total reads | Daphnia pulex | Ankistrodesmus falcatus | Raphidocelis subcapitata | Scenedesmus quadricauda | Brachionus plicatilis | Volvox cateri | Stylonychia lemnae | Chlamydomonas eustigma |
|---|---|---|---|---|---|---|---|---|---|
| eRNA_20T1 | 194,229,708 | 773,529 | 5,105,899 | 13,745,980 | 14,343,163 | 5,755,453 | 9,482,234 | 12,912,193 | 20,673,456 |
| eRNA_20T2 | 164,328,302 | 1,056,794 | 4,158,454 | 9,423,422 | 10,577,585 | 4,326,777 | 6,733,253 | 7,792,666 | 15,569,419 |
| eRNA_20T3 | 166,935,050 | 2,007,126 | 4,259,204 | 12,076,877 | 12,731,338 | 5,300,567 | 9,960,792 | 8,768,980 | 20,287,504 |
| eRNA_20T4 | 111,657,602 | 963,726 | 2,203,949 | 7,615,700 | 8,420,941 | 3,201,828 | 5,569,258 | 6,202,984 | 12,431,769 |
| eRNA_28T1 | 158,851,649 | 556,462 | 2,065,240 | 6,826,188 | 7,207,397 | 4,286,280 | 4,563,917 | 7,100,241 | 8,015,263 |
| eRNA_28T2 | 161,632,735 | 115,806 | 2,041,879 | 5,564,659 | 6,050,008 | 3,503,434 | 4,132,142 | 6,192,668 | 7,719,423 |
| eRNA_28T3 | 142,601,838 | 165,286 | 3,864,896 | 7,606,623 | 8,099,620 | 3,574,020 | 5,205,967 | 8,316,693 | 9,469,856 |
| eRNA_28T4 | 169,831,709 | 746,323 | 3,187,047 | 12,499,495 | 13,589,887 | 3,556,469 | 8,894,525 | 11,025,956 | 18,645,765 |
| eRNA_avg | 158,758,574 | 798,132 | 3,360,821 | 9,419,868 | 10,127,492 | 4,188,104 | 6,817,761 | 8,539,048 | 14,101,557 |
| eRNA_20avg | 159,287,666 | 1,200,294 | 3,931,877 | 10,715,495 | 11,518,257 | 4,646,156 | 7,936,384 | 8,919,206 | 17,240,537 |
| eRNA_28avg | 158,229,483 | 395,969 | 2,789,766 | 8,124,241 | 8,736,728 | 3,730,051 | 5,699,138 | 8,158,890 | 10,962,577 |
| oRNA_20T1 | 267,407,877 | 244,968,635 | NA | NA | NA | NA | NA | NA | NA |
| oRNA_20T2 | 262,069,830 | 246,003,499 | NA | NA | NA | NA | NA | NA | NA |
| oRNA_20T3 | 212,740,522 | 198,481,152 | NA | NA | NA | NA | NA | NA | NA |
| oRNA_20T4 | 218,895,465 | 207,005,534 | NA | NA | NA | NA | NA | NA | NA |
| oRNA_28T1 | 184,893,400 | 175,093,620 | NA | NA | NA | NA | NA | NA | NA |
| oRNA_28T2 | 186,039,156 | 178,254,098 | NA | NA | NA | NA | NA | NA | NA |
| oRNA_28T3 | 154,638,938 | 146,305,180 | NA | NA | NA | NA | NA | NA | NA |
| oRNA_28T4 | 203,208,790 | 186,591,250 | NA | NA | NA | NA | NA | NA | NA |
| oRNA_avg | 211,236,747 | 197,837,871 | NA | NA | NA | NA | NA | NA | NA |
| oRNA_20avg | 240,278,424 | 224,114,705 | NA | NA | NA | NA | NA | NA | NA |
| oRNA_28avg | 182,195,071 | 171,561,037 | NA | NA | NA | NA | NA | NA | NA |

**Table S3**. The number of genes detected and identified in eRNA samples as differentially expressed for the four eukaryotes (*D. pulex, A. falcatus, R. subcapitata* and *S. quadricauda*) that were a priori known to persist in the tanks and the five most highly abundant species identified by Diamond alignment against the GenBank nr database. A false discovery rate adjusted p-value <0.05 was used to determine significantly differentially expressed genes.

| Species | # genes detected | # differentially expressed genes |
|---|---|---|
| *Daphnia pulex* | 3,919 | 32 |
| *Ankistrodesmus falcatus* | 1,982 | 44 |
| *Raphidocelis subcapitata* | 259 | 12 |
| *Scenedesmus quadricauda* | 3,180 | 20 |
| *Brachionus plicatilis* | 144 | 0 |
| *Volvox cateri* | 264 | 4 |
| *Stylonychia lemnae* | 99 | 7 |
| *Chlamydomonas eustigma* | 179 | 2 |

**Table S4.** Statistics of KEGG KO IDs differentially expressed (FDR-adjusted p-value <0.05) between 20°C and 28°C eRNA samples.

| Name | Entry | Symbol | log2Fold Change | FDR-adjusted p-value |
|---|---|---|---|---|
| malate-CoA ligase subunit beta [EC:6.2.1.9] | K14067 | mtkA | -12.47 | 1.49E-23 |
| malate-CoA ligase subunit alpha [EC:6.2.1.9] | K08692 | mtkB | -12.33 | 1.07E-22 |
| cyclin-dependent kinase 4 [EC:2.7.11.22] | K02089 | CDK4 | -11.20 | 3.42E-16 |
| prophage regulatory protein | K07733 | alpA | 11.25 | 3.37E-13 |
| heat-stable enterotoxin receptor [EC:4.6.1.2] | K12320 | GUCY2C | -24.62 | 3.51E-13 |
| solute carrier family 25, member 43 | K15120 | SLC25A43 | 26.69 | 5.29E-12 |
| spore germination protein | K06306 | yaaH | 23.97 | 5.48E-12 |
| mycobactin phenyloxazoline synthetase | K04788 | mbtB | 23.77 | 1.45E-11 |
| DNA-directed RNA polymerase III subunit RPC7 | K03024 | RPC7, POLR3G | 23.53 | 2.10E-11 |
| disintegrin and metalloproteinase domain-containing protein 18 [EC:3.4.24.-] | K16909 | ADAM18 | 23.61 | 4.06E-11 |
| conjugal transfer pilus assembly protein TraA | K12069 | traA | -10.73 | 4.06E-11 |
| hemicentin | K17341 | HMCN | 4.99 | 1.01E-09 |
| acyl-CoA (8-3)-desaturase (Delta-5 desaturase) [EC:1.14.19.44] | K10224 | FADS1 | 23.18 | 2.91E-09 |
| annexin A6 | K17094 | ANXA6 | 3.16 | 2.18E-08 |
| GTP pyrophosphokinase [EC:2.7.6.5] | K07816 | E2.7.6.5 | -10.02 | 7.87E-07 |
| P-type Na+/K+ transporter [EC:7.2.2.3 7.2.2.-] | K01536 | ENA | 7.51 | 1.94E-06 |
| violacein biosynthesis protein VioB | K20087 | vioB | 10.16 | 3.19E-06 |
| lantibiotic biosynthesis protein | K20483 | nisB, spaB, epiB | -9.81 | 1.07E-05 |
| putative tricarboxylic transport membrane protein | K07795 | tctC | -2.08 | 1.31E-05 |
| sn-glycerol 3-phosphate transport system substrate-binding protein | K05813 | ugpB | -2.60 | 2.06E-05 |
| aqualysin 1 [EC:3.4.21.111] | K20754 | pstI | 3.09 | 3.70E-05 |
| calpain-2 [EC:3.4.22.53] | K03853 | CAPN2 | 9.49 | 4.40E-05 |
| midline 1 [EC:2.3.2.27] | K08285 | TRIM18, MID1 | 11.24 | 5.84E-05 |
| glyoxylate/hydroxypyruvate reductase [EC:1.1.1.79 1.1.1.81] | K15919 | HPR2_3 | -9.07 | 1.15E-04 |
| potassium channel subfamily K, other eukaryote | K05389 | KCNKF | -3.25 | 2.03E-04 |
| octopine/nopaline transport system substrate-binding protein | K10018 | occT, nocT | -8.87 | 3.33E-04 |
| candicidin polyketide synthase FscC | K20787 | fscC | 10.58 | 9.29E-04 |
| methyl acetate hydrolase [EC:3.1.1.114] | K18372 | acmB | -3.11 | 1.22E-03 |
| betaine-homocysteine S-methyltransferase [EC:2.1.1.5] | K00544 | BHMT | 9.04 | 1.22E-03 |
| katanin p60 ATPase-containing subunit A1 [EC:5.6.1.1] | K07767 | KATNA1 | 2.31 | 1.54E-03 |
| 5,10-methylenetetrahydromethanopterin reductase [EC:1.5.98.2] | K00320 | mer | -8.86 | 1.54E-03 |
| gem associated protein 5 | K13133 | GEMIN5 | -8.62 | 1.79E-03 |
| glucose/mannose transport system permease protein | K17317 | gtsC, glcG | -8.60 | 1.79E-03 |
| NAD(P)H dehydrogenase (quinone) [EC:1.6.5.2] | K19267 | qorB | 2.62 | 1.79E-03 |
| 3,4-dihydroxy 2-butanone 4-phosphate synthase [EC:4.1.99.12] | K02858 | ribB, RIB3 | 10.72 | 1.98E-03 |

| | | | | |
|---|---|---|---|---|
| 3,4-dihydroxy-9,10-secoandrosta-1,3,5(10)-triene-9,17-dione 4,5-dioxygenase [EC:1.13.11.25] | K16049 | hsaC | -8.42 | 2.10E-03 |
| phosphatidylinositol phospholipase C, gamma-2 [EC:3.1.4.11] | K05859 | PLCG2 | -10.35 | 2.19E-03 |
| uncharacterized protein | K06884 | K06884 | -8.62 | 2.21E-03 |
| cathepsin F [EC:3.4.22.41] | K01373 | CTSF | 2.19 | 2.68E-03 |
| rhamnogalacturonan endolyase [EC:4.2.2.23] | K18195 | RGL4, rhiE | 2.68 | 3.34E-03 |
| two-component system, OmpR family, response regulator ResD | K07775 | resD | 8.86 | 3.34E-03 |
| fructose transport system substrate-binding protein | K10552 | frcB | -1.40 | 3.94E-03 |
| methionine transaminase [EC:2.6.1.88] | K14287 | ybdL | -8.46 | 4.85E-03 |
| mitogen-activated protein kinase kinase 2 [EC:2.7.12.2] | K20603 | MKK2 | 6.90 | 4.85E-03 |
| uncharacterized protein | K09794 | K09794 | -8.30 | 5.58E-03 |
| trans-aconitate 2-methyltransferase [EC:2.1.1.144] | K00598 | tam | 2.86 | 5.84E-03 |
| mycothiol S-conjugate amidase [EC:3.5.1.115] | K18455 | mca | -9.92 | 5.84E-03 |
| osomolarity two-component system, sensor histidine kinase NIK1 [EC:2.7.13.3] | K19691 | NIK1, TCSC | 3.14 | 6.23E-03 |
| glutamate receptor 3 | K05199 | GRIA3 | -8.26 | 6.39E-03 |
| putative tricarboxylic transport membrane protein | K07793 | tctA | -2.22 | 8.11E-03 |
| mitogen-activated protein kinase 3 [EC:2.7.11.24] | K20536 | MPK3 | 2.73 | 8.90E-03 |
| putative spermidine/putrescine transport system substrate-binding protein | K02055 | ABC.SP.S | -1.65 | 1.02E-02 |
| oligopeptide transport system substrate-binding protein | K15580 | oppA, mppA | -2.09 | 1.07E-02 |
| chlorophyllide a oxygenase [EC:1.14.13.122] | K13600 | CAO | -4.78 | 1.14E-02 |
| solute carrier family 24 (sodium/potassium/calcium exchanger), member 2 | K13750 | SLC24A2, NCKX2 | 12.06 | 1.29E-02 |
| alpha-1,3-mannosyltransferase [EC:2.4.1.-] | K13690 | CMT1 | -9.78 | 1.31E-02 |
| E3 ubiquitin-protein ligase HERC1 [EC:2.3.2.26] | K10594 | HERC1 | -4.28 | 1.31E-02 |
| fimbrial chaperone protein | K07346 | fimC | 8.17 | 1.31E-02 |
| stage II sporulation protein M | K06384 | spoIIM | -9.50 | 1.36E-02 |
| demethylspheroidene O-methyltransferase [EC:2.1.1.210] | K09846 | crtF | 8.38 | 1.39E-02 |
| divalent anion:Na+ symporter, DASS family | K03319 | TC.DASS | -1.75 | 1.41E-02 |
| L-galactono-1,4-lactone dehydrogenase [EC:1.3.2.3] | K00225 | GLDH | -1.38 | 1.47E-02 |
| ADP-L-glycero-D-manno-heptose 6-epimerase [EC:5.1.3.20] | K03274 | gmhD, rfaD | -2.71 | 1.47E-02 |
| heat shock transcription factor 3 | K09416 | HSF3 | 5.95 | 1.47E-02 |
| homeobox protein homothorax | K16672 | HTH | 9.56 | 1.52E-02 |
| oxidoreductase [EC:1.1.1.-] | K16015 | rifL, asm44 | -9.52 | 1.53E-02 |
| uncharacterized protein | K09857 | K09857 | 8.24 | 1.57E-02 |
| trafficking protein particle complex subunit 9 | K20306 | TRAPPC9, TRS120 | -9.52 | 1.64E-02 |
| beta-glucosidase [EC:3.2.1.21] | K01188 | E3.2.1.21 | 4.08 | 1.64E-02 |
| uncharacterized protein | K07161 | K07161 | -7.98 | 2.00E-02 |
| isoflavone/4'-methoxyisoflavone 2'-hydroxylase [EC:1.14.14.90 1.14.14.89] | K13260 | CYP81E | 9.71 | 2.29E-02 |
| major outer membrane protein P.IB | K18133 | K18133, porB | -9.31 | 2.32E-02 |
| 4-hydroxy-2-oxovalerate/4-hydroxy-2-oxohexanoate aldolase [EC:4.1.3.39 4.1.3.43] | K18365 | bphI, xylK, nahM, tesG | -6.59 | 2.45E-02 |
| cellular nucleic acid-binding protein | K09250 | CNBP | 3.95 | 2.93E-02 |
| prolactin regulatory element-binding protein | K14003 | PREB, SEC12 | 8.17 | 2.96E-02 |

| | | | | |
|---|---|---|---|---|
| dual oxidase maturation factor 1 | K17233 | DUOXA1 | -9.28 | 2.96E-02 |
| arylsulfatase I/J [EC:3.1.6.-] | K12375 | ARSI_J | 1.23 | 3.22E-02 |
| TRAP-type transport system periplasmic protein | K21395 | yiaO | -2.02 | 3.27E-02 |
| midasin | K14572 | MDN1, REA1 | 1.08 | 3.82E-02 |
| aquaporin SIP | K09875 | SIP | -9.20 | 3.88E-02 |
| methyl-accepting chemotaxis protein WspA | K13487 | wspA | -1.76 | 3.95E-02 |
| peptidase inhibitor 16 | K20412 | PI16, CRISP9, CD364 | 2.44 | 4.03E-02 |
| NitT/TauT family transport system substrate-binding protein | K02051 | ABC.SN.S | -1.66 | 4.03E-02 |
| nuclear pore complex protein Nup85 | K14304 | NUP85 | -9.17 | 4.03E-02 |
| cell division protein ZapE | K06916 | zapE | -9.11 | 4.47E-02 |
| butyrate response factor | K18753 | ZFP36L | 1.73 | 4.47E-02 |
| sphingolipid 8-(E/Z)-desaturase [EC:1.14.19.29] | K21734 | SLD | 6.10 | 4.47E-02 |
| molybdopterin synthase sulfur carrier subunit | K21232 | MOCS2A, CNXG | 7.90 | 4.47E-02 |
| calpain-12 [EC:3.4.22.-] | K04740 | CAPN12 | 1.79 | 4.73E-02 |
| atrial natriuretic peptide clearance receptor | K12325 | NPR3 | -7.73 | 4.73E-02 |
| butyrophilin | K06712 | BTN, CD277 | 4.94 | 4.73E-02 |
| E3 ubiquitin-protein ligase RGLG [EC:2.3.2.27] | K16280 | RGLG | 1.57 | 4.73E-02 |
| betaine lipid synthase | K13621 | BTA1 | -1.50 | 4.97E-02 |

**Table S5.** Statistics for the 4/62 *Daphnia pulex* heat shock proteins identified as significantly but lowly upregulated (log 2 fold change > 1) in organismal RNA. The statistics for environmental RNA (eRNA) are shown on the right. NA indicates that dp_genes 3128 and 13761 were not detected in eRNA samples, and that FDR adjusted p-value could not be calculated for dp_gene 10006. Within RNA type, a gene was considered to be detected if it had sum ≥ 10 counts in either temperature conditions. Differential gene expression analyses were conducted using DESeq2.

| | Organismal RNA | | Environmental RNA | |
|---|---|---|---|---|
| Gene | log2 Fold Change | FDR adjusted p-value | log2 Fold Change | FDR adjusted p-value |
| dp_gene3128 | 1.35 | 9.01e-14 | NA | NA |
| dp_gene9537 | 1.20 | 1.88e-6 | -0.29 | 0.95 |
| dp_gene10006 | 1.21 | 3.37e-5 | 4.10 | NA |
| dp_gene13761 | 1.12 | 0.004 | NA | NA |
